# Large-scale remote sensing reveals that tree mortality in Germany appears to be greater than previously expected

**DOI:** 10.1101/2024.11.10.622853

**Authors:** Felix Schiefer, Sebastian Schmidtlein, Henrik Hartmann, Florian Schnabel, Teja Kattenborn

## Abstract

Global warming poses a major threat to forests and events of increased tree mortality are observed globally. Studying tree mortality often relies on local-level observations of dieback while large-scale analyses are lacking. Satellite remote sensing provides the spatial coverage and sufficiently high temporal and spatial resolution needed to investigate tree mortality at landscape-scale. However, adequate reference data for training satellite-based models are scarce. In this study, we employed the first maps of standing deadwood in Germany for the years 2018–2022 with 10 m spatial resolution that were created by using tree mortality observations spotted in hundreds of drone images as the reference. We use these maps to study spatial and temporal patterns of tree mortality in Germany and analyse their biotic and abiotic environmental drivers using random forest regression. In 2019, the second consecutive hotter drought year in a row, standing deadwood increased steeply to 334 ± 189 kilohectar (kha) which corresponds to 2.5 ± 1.4% of the total forested area in Germany. *Picea abies*, *Pinus sylvestris*, and *Fagus sylvatica* showed highest shares of standing deadwood. During 2018–2021 978 ± 529 kha (7.9 ± 4.4%) of standing dead trees accumulated. The higher mortality estimates that we report compared to other surveys (such as the ground-based forest condition survey) can be partially attributed to the fact that remote sensing captures mortality from a bird’s eye perspective and that the high spatial detail (10 m) in this study also captures scattered occurrences of tree mortality. Atmospheric drought (i.e., climatic water balance and vapor pressure deficit) and temperature extremes (i.e., number of hot days and frosts after vegetation onset) were the most important predictors of tree mortality. We found increased tree mortality for smaller and younger stands and also on less productive sites. Monospecific stands were generally not more affected by mortality, but only when interactions with damaging insects (e.g., bark beetles) occurred. Because excess tree mortality rates threaten many forests across the globe, similar analyses of tree mortality are warranted and technically feasible at the global scale. We encourage the international scientific community to share and compile local data on deadwood occurrences (see example: www.deadtrees.earth) as such a collaborative effort is required to help understand mortality events on a global scale.

## Introduction

Forests cover one third of the total land surface (FAO, 2020), are an important carbon sink, and provide a range of ecosystem services. Climate change and the associated rise in temperatures, the occurrence of episodic precipitation and droughts, or an increased atmospheric water vapor deficit pose a major threat to forests (Hartmann et al., 2022; McDowell et al., 2022; Schuldt et al., 2020). Consequently, trees get stressed and may eventually die due to carbon starvation, hydraulic failure, or ensuing pest infestations. This cascade of effects is particularly pronounced during ‘hotter droughts’, in which long periods of drought coincide with high temperatures (Allen et al., 2015; Hammond et al., 2022). With the rise in average temperatures, an earlier start to the growing season has also been observed in recent years. Nevertheless, late-frost events can still occur after bud burst, which can weaken the trees, prevent resprouting, and may ultimately lead to mortality (Vanoni et al., 2016).

We currently witness the emergence of hotter droughts even in temperate regions (Hammond et al., 2022; Hartmann et al., 2022). Consecutive (hotter) droughts have also become more frequent (Hari et al., 2020; Rakovec et al., 2022) and large-scale tree stress responses and diebacks have been observed after the prolonged 2018–2021 drought in Central Europe (Rakovec et al., 2022; Schnabel et al., 2022; Schuldt et al., 2020) and the 2012–2016 drought in Northern America (Byer & Jin, 2017). Even if the trees do not die in a first year of drought critical ecosystem changes and mortality may still occur in the years after the drought, known as drought legacy or lag effects (Obladen et al., 2021; Pohl et al., 2023; Schnabel et al., 2022). Due to a climate that is increasingly characterized by extremes, all these stress responses can also occur together, creating compound effects (Zscheischler et al., 2018). Understanding the drivers that lead to tree mortality is crucial to assess and project the impact of ongoing global warming on forests and to adapt forest management strategies accordingly.

Drivers and dynamics of tree mortality have been assessed in local studies but the findings can often not be generalized to larger scales (Clark et al., 2016). Standardized surveys, such as the national forest condition survey in Germany (BMEL, 2023), provide a countrywide estimate of forest health and tree mortality. However, such initiatives usually feature small sample sizes that—given large environmental and forest structural variability—do not suffice to robustly understand spatiotemporal tree mortality dynamics at the landscape-level. Consequently, suitable large-scale and representative data sets on tree mortality are lacking, and many of our findings on the drivers of tree mortality arise from compiled and harmonized data sets of coarse *in situ* observations of dieback events (Allen et al., 2010; Hammond et al., 2022). This poses two problems for the investigation of the underlying causes: large-scale occurrences of standing deadwood often only accumulate over time. The temporal link between the environmental cause and the dieback event may therefore be weakened or already overlaid by other factors. Furthermore, scattered and gradual occurrences of tree mortality are likely to be underrepresented in the aforementioned data sets. (Cheng et al., 2024; Milodowski et al., 2017).

The lack of large-scale tree mortality data constrains our understanding on the roles of abiotic and biotic environmental drivers. Climate change and associated pest and pathogen outbreaks have increased the risk of large-scale tree mortality (Allen et al., 2010; Huang et al., 2020; McDowell et al., 2020), but spatiotemporal dependencies often remain unknown. For example, Socha et al. (2023) suggest that higher productivity and greater tree age enhance susceptibility to drought-induced mortality. While smaller and younger trees die mainly due to competition for resources (Kulha et al., 2023; Stephenson & Das, 2020), larger and older trees have an accumulated risk of disease and damage (Bennett et al., 2015; Stovall et al., 2019, 2020). The influence of tree species richness on mortality remains ambiguous as well (Depauw et al., 2024). While a positive effect of tree species richness on the stability and productivity of forests is assumed to be likely due to performance enhancing and buffering effects of diversity (Anderegg et al., 2018; Schnabel et al., 2021), studies found mixed results regarding the effect of tree diversity on forest responses to drought (Grossiord, 2020). While recent experimental work points towards a slightly positive or non-significant effect of tree diversity on individual tree or stand level tree mortality (Blondeel et al., 2024; Liu et al., 2022), Searle et al. (2022) found higher tree diversity to be correlated with higher stand level tree mortality using forest inventory plots from across Canada and the United States. These at a first glance contradictory results clearly illustrate that tree mortality is still not generically understood. The relationships that have been identified at the individual tree level have not been investigated or confirmed at the landscape level and require further research (McDowell et al., 2022). This is particularly important as tree species differ in their response to changes in biotic and abiotic conditions, yet large-scale studies only rarely differentiate the effects of the various environmental drivers at the species level (Wegler & Kuenzer, 2024). Thus, our understanding of tree mortality dynamics across gradients of forest composition, forest structure, and environmental conditions requires data on tree mortality dynamics at large scale and sufficient spatial detail. Large-scale patterns of tree mortality can be revealed using Earth observation satellite missions (Brodrick & Asner, 2017; Byer & Jin, 2017; Garrity et al., 2013; Hansen et al., 2013; Schwantes et al., 2016). Previous attempts for large-scale mapping of tree mortality are often based on satellite data that usually feature spatial resolutions that are much coarser than the tree canopies being targeted, such as MODIS with 250 m or Landsat with 30 m resolution (Byer & Jin, 2017; Campbell et al., 2020; Schuldt et al., 2020; Schwantes et al., 2016). As such coarse spatial resolutions cannot adequately reveal small-scaled or scattered tree mortality (Hartmann, Moura, et al., 2018), a series of studies used higher resolution data, for example 10 m spatial resolution of Sentinel-2 (Thonfeld et al., 2022), 3 m of PlanetScope (Francini et al., 2020), or even down to 0.5 m of WorldView and Quickbird (Garrity et al., 2013; Liu et al., 2021). However, commonly such data is not being used to map tree mortality directly because reference data on tree mortality is missing for training supervised models (Eliades et al., 2024). Instead, most studies use vegetation indices as a proxy of dieback. The causes for such changes in vegetation indices are manifold and may not necessarily be linked to tree mortality but rather to an overall vitality or health decline. Therefore, Schiefer et al. (2023) presented an approach that explicitly maps tree mortality using hundreds of drone images as training data for supervised models. This approach builds on the fact that standing dead trees can be precisely and automatically segmented in drone images (local scale), which provide an effective source to generate large samples of training data for predicting tree mortality with Sentinel satellite data at large scales (see Figure 1). Their upscaling approach resulted in Germany-wide maps of fractional cover of standing deadwood at the original Sentinel image resolution of 10 m for the consecutive years 2018–2021. Here, we use this dataset to reveal the spatiotemporal dynamics of standing deadwood in Germany, and studied the species-specific patterns using tree species information from Blickensförfer et al. (2024). We further investigate the influence of several biotic and abiotic environmental drivers on the observed tree mortality at landscape level.

**Figure 1.**
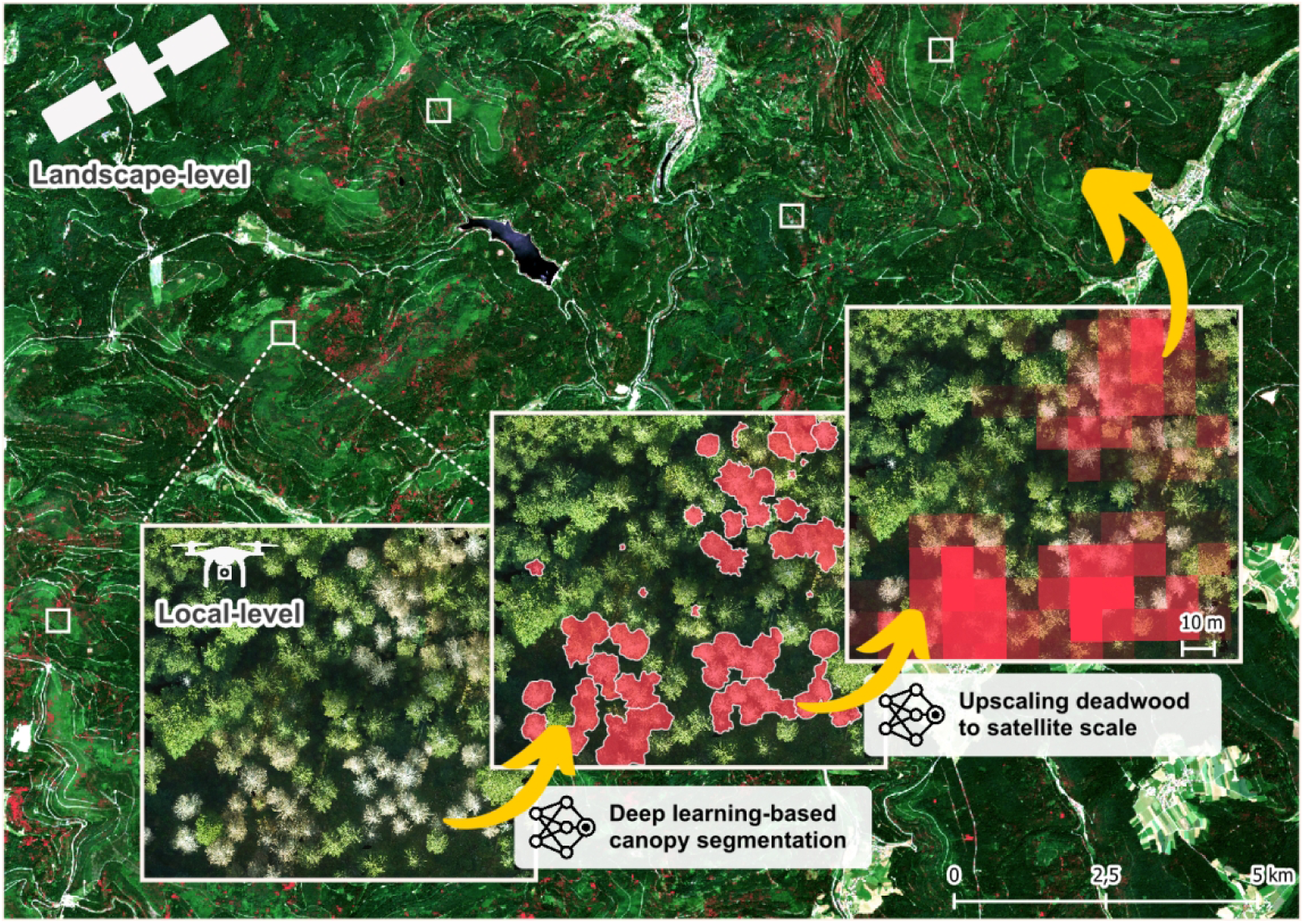
Schematic overview of the presented upscaling approach in Schiefer et al. (2023) extrapolating local observations of standing deadwood in UAV-imagery to landscape-level predictions based on satellite time series data.

## Methods

### Tree mortality in Germany

To identify and study the pattern of tree mortality, we extracted fractional cover values of standing dead trees for the available years 2018–2021 from the Germany-wide layers from Schiefer et al. (2023), with an interactive preview available at https://deadtrees.earth/. Schiefer et al. (2023) automatically segmented crowns of standing dead trees from very high-resolution drone imagery available at local scales using Convolutional Neural Networks (CNN). They converted these segmentations to cover fractions of standing deadwood per 10 m Sentinel-2 pixel and fed them into long short-term memory networks (LSTM) for large-scale extrapolation using time series of Sentinel-1 and Sentinel-2 as predictors. The final Germany-wide maps show the cover fraction [%] of standing deadwood in the tree canopy for a given satellite pixel at 10 m spatial resolution (RMSE = 22%).

As standing deadwood is not always necessarily removed from the forest, it can appear in a pixel in consecutive years. Therefore, we only considered the year of first occurrence of standing deadwood. This ensured an explicit link between occurrences of standing deadwood and the prevailing environmental conditions. We defined the year of the first occurrence as the point in time at which the fractional cover exceeds a certain threshold for the first time. Only this year was then considered in the subsequent analysis. To avoid bias in the results by setting an arbitrary threshold, we applied different thresholds of standing deadwood continuously between 30% and 70%. We then averaged the results of the different thresholds in a sensitivity analysis and calculated standard deviations to provide uncertainty estimates of the detected standing deadwood. The yearly binary maps of first occurrence then only served as a mask and in subsequent analysis again the percentages of standing deadwood were used. We then overlaid the standing deadwood maps with maps of the dominant tree species by Blickensdörfer et al. (2024). The species maps are based on national forest inventory data and remote sensing and depict the spatial distribution of 11 tree species in Germany, i.e., *Picea abies* (31.4% area share, 85.57 ± 0.12 User’s accuracy), *Pinus sylvestris* (22%, 94.96 ± 0.10), *Fagus sylvatica* (15,1%, 75.44 ± 0.12), *Quercus spp.* (8,9%, 57.65 ± 0.18), *Larix spp.* (2.6%, 47.29 ± 0.53), *Alnus glutinosa* (2.6%, 64.67 ± 0.53), *Betula pendula* (2,1%, 46.76 ± 0.93), *Pseudotsuga menziesii* (2%, 39.46 ± 0.35), *Abies alba* (2%, 64.29 ± 0.41), and two other deciduous species groups. The map of Blickensdörfer et al. (2024) has a spatial resolution of 10 m.

We assessed regional differences for Germany’s 73 major natural regions first defined by Meynen and Schmithüsen (1953) adapted by the Federal Office for Nature Conservation (Bundesamt für Naturschutz–BfN). The major natural regions divide Germany into physical units of similar geography primarily based on geomorphological, geological, hydrological, and pedological criteria. They allow for a regional comparability beyond administrative boundaries.

### Identifying predictors of tree mortality using random forest

To identify the main predictors of tree mortality in Germany, we calculated random forest models (Breiman, 2001) using the previously described standing deadwood maps as a response and 21 environmental predictor variables (Table 1). To rule out multicollinearity among the predictors, we dropped variables with correlations higher than 0.7, as this threshold is considered an appropriate rule of thumb for critical levels of collinearity (Dormann et al., 2013), so that the ecologically more meaningful variable remained (Figure 2). An overview of the originally selected environmental variables along with the rational for the selection of variables can be found in the Appendix.

**Table 1.** Environmental predictor variables.

| Variable | Spatial resolution | Unit | Rationale | Data source |
| --- | --- | --- | --- | --- |
| Slope | 10 m | ° | Topographic site conditions | Slope and eastness (sinus) and northness (cosine) of terrain aspect derived from Copernicus DEM EEA-10 (2021) |
| Eastness |  | rad |  |  |
| Northness |  | rad |  |  |
| Sand content | 30 m | % | Soil type | Hengl, T. and Parente, L. (2022) |
| Stand age | 100 m | years | Older trees might be more susceptible to mortality (cf. Socha et al., 2023) | Besnard et al. (2021) |
| Canopy height | 10 m | m | Taller trees might be more susceptible to mortality (cf. Bennett et al., 2015; Stovall et al., 2019, 2020) | Lang et al. (2023) |
| Tree species richness | 10 m | no. | Tree species richness might mitigate mortality (cf. Anderegg et al., 2018; Depauw et al., 2024; Grossiord, 2020; Schnabel et al., 2021; Searle et al., 2022) | Calculated from Blickensdörfer et al. (2024) as number of tree species in a 100 m buffer |
| Temperature annual range | 1 km | °C | Climatic site conditions | Derived from Fick & Hijmans (2017) |
| Precipitation seasonality |  | mm |  |  |
| Precipitation of warmest quarter |  | mm |  |  |
| Late frosts* | 1 km | days | Frost damage after bud burst (cf. Vanoni et al., 2016) | Calculated from vegetation onset (DWD, 2023b) and CERRA reanalysis (Schimanke et al., 2021) |
| Soil drought intensity* | 4 km | - | Edaphic site conditions | During vegetation period (Zink et al., 2016) |
| Vapor pressure deficit | 5.5 km | Pa | Decreasing water availability increases mortality risk (Allen et al., 2010) | Calculated as 75%-quartile of daily maximum during growing period (March-October) from CERRA reanalysis (Schimanke et al., 2021) |
| Climatic water balance* | 1 km | mm |  | Calculated from precipitation and potential evapotranspiration over grass (DWD, 2023d, 2023e) as anomaly from reference period 1961-1990 (DWD, 2023f) |
| Hot days* | 1 km | days | Heat-induced tree mortality through cavitation of water columns within the xylem (Allen et al., 2010) | (DWD, 2023a) |
| Biomass | 100 m | Mg/ha | Proxy for site productivity. Hypothesis that higher productivity enhances susceptibility to drought-induced mortality (cf. Socha et al., 2023) | ESA Biomass CCI (Santoro & Cartus, 2023) |
| Net Primary Productivity | 500 m | kgC/m<sup>2</sup>/year |  | Running, S. and Zhao, M. (2021) |
| *Variable also for the year prior to first occurrence of standing deadwood |  |  |  |  |

**Figure 2.**
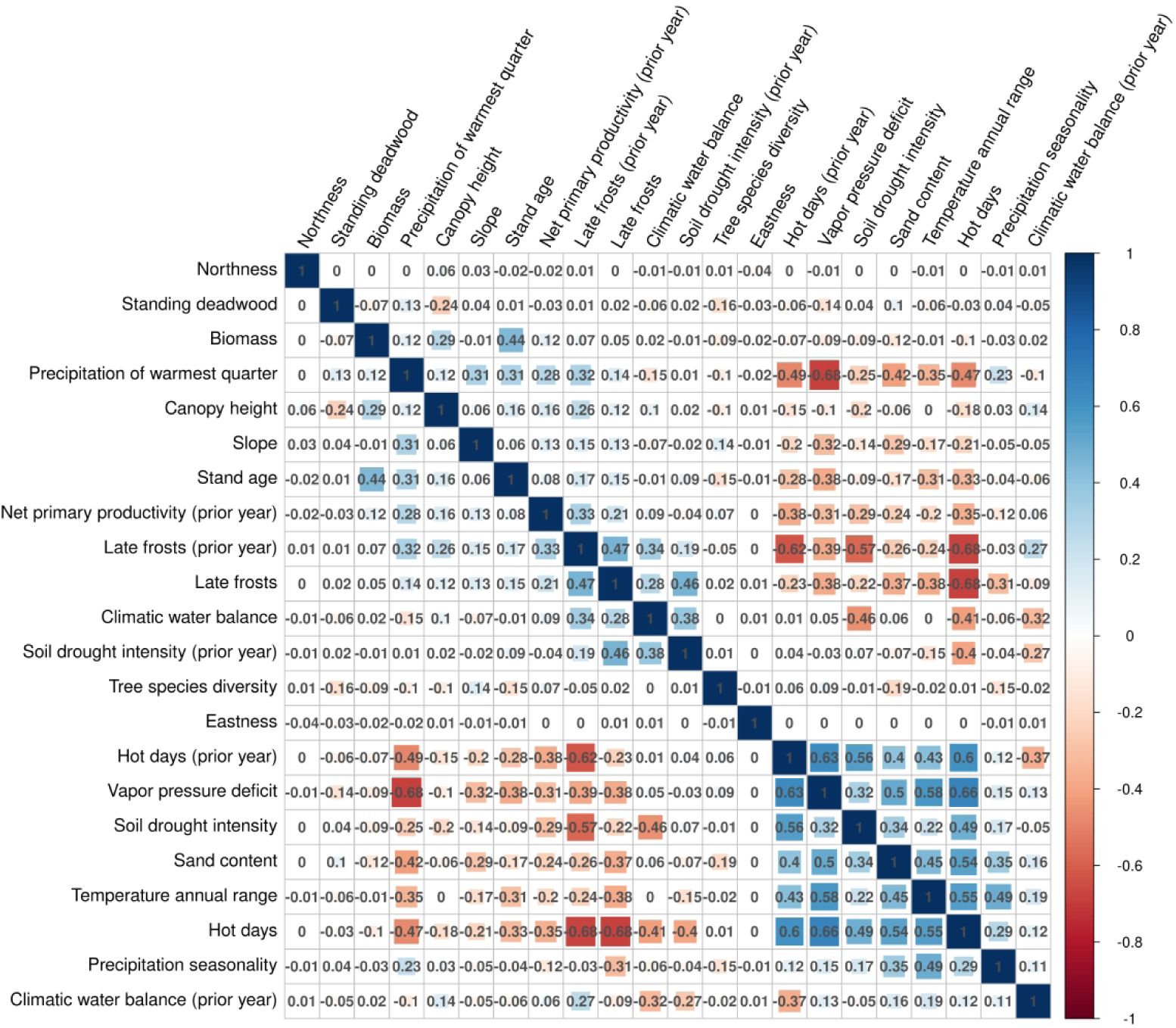
Correlation plot for the selected 21 environmental predictor variables and standing deadwood. Variables with correlations higher than 0.7 were dropped for subsequent analysis.

We separately analysed the seven main tree species—as defined by the German national forest inventory—*Fagus sylvatica*, *Quercus spp.*, *Picea abies*, *Pinus sylvestris*, *Abies alba*, *Pseudotsuga menziesii*, and *Larix spp.*. We ran random forest models for each species individually. For every model, we randomly sampled 32,000 pixels, half with and half without standing deadwood. To build temporally generalising models, we sampled 4,000 pixels per each year of first occurrence. For every model, we calculated permutation importances and accumulated local effects (ALE, Apley & Zhu (2020). Permutation importance measures the increase in model prediction error after permuting a variable’s values and is a straightforward means for identifying important variables. ALE further reveal how a variable influences model predictions on average and can help understand the direction of a variable’s effect. ALE is an alternative to the frequently applied partial dependence methods (Friedman, 2001) but they are less prone to erroneous results due to intercorrelated variables and are less computational demanding (Apley & Zhu, 2020). We repeated this procedure 100 times for every species and averaged the permutation importances and accumulated local effects. We calculated *Moran’s I* based on the model residuals. All values were close to 0 (± 0.04) and we could therefore rule out spatial autocorrelation. All analyses were conducted in R language (R Core Team, 2022) using the packages *ranger* for the random forest implementation (Wright & Ziegler, 2017) and *iml* for the accumulated local effects (Molnar et al., 2018).

## Results

In 2018 1.4 ± 1.0% and an approximate area of 179 ± 126 kilohectares (kha) of Germany’s forests were dead (Figure 3). The largest share corresponds to *Pinus* with 113 ± 81 kha, which equates to 4.4 ± 2.9% of all *Pinus* trees in Germany. In 2019 standing deadwood increased for most species and particularly for *Picea* (6.3 ± 2.1%, 88 ± 39 kha) and *Pinus* (6.7 ± 3.8%, 201 ± 118 kha). 334 ± 189 kha of forest died in 2019, approximately 2.5 ± 1.4% of the forested area in Germany. In 2020 the total amount of new standing deadwood slightly decreased to 2.6 ± 1.3% and 307 ± 151 kha. For *Picea* (8.3 ± 2.5%, 146 ± 44 kha), *Fagus* (3.5 ± 1.5%, 17 ± 8.2 kha) and *Quercus* (2.1 ± 0.8%, 2.1 ± 1.2 kha) standing deadwood peaked in 2020. In 2021, mortality rates further decreased for all species. Accumulated over the years, in total 978 ± 529 kha of trees died from 2018 to 2021, accounting for 7.9 ± 4.4% of the forested area in Germany. For the four most frequent tree species in Germany, the accumulated standing deadwood from 2018 to 2021 was: *Fagus* 8.2 ± 3.8%, *Picea* 21.5 ± 7.1, *Pinus* 20 ± 12%, and *Quercus* 4.5 ± 1.7%. The variation around the mean of the different threshold values for the determination of the first occurrence of standing deadwood was high for some species, particularly *Pinus.* Detailed statistics of the temporal development of standing deadwood for the different species can be found in the supplementary materials (Table S2).

**Figure 3.**
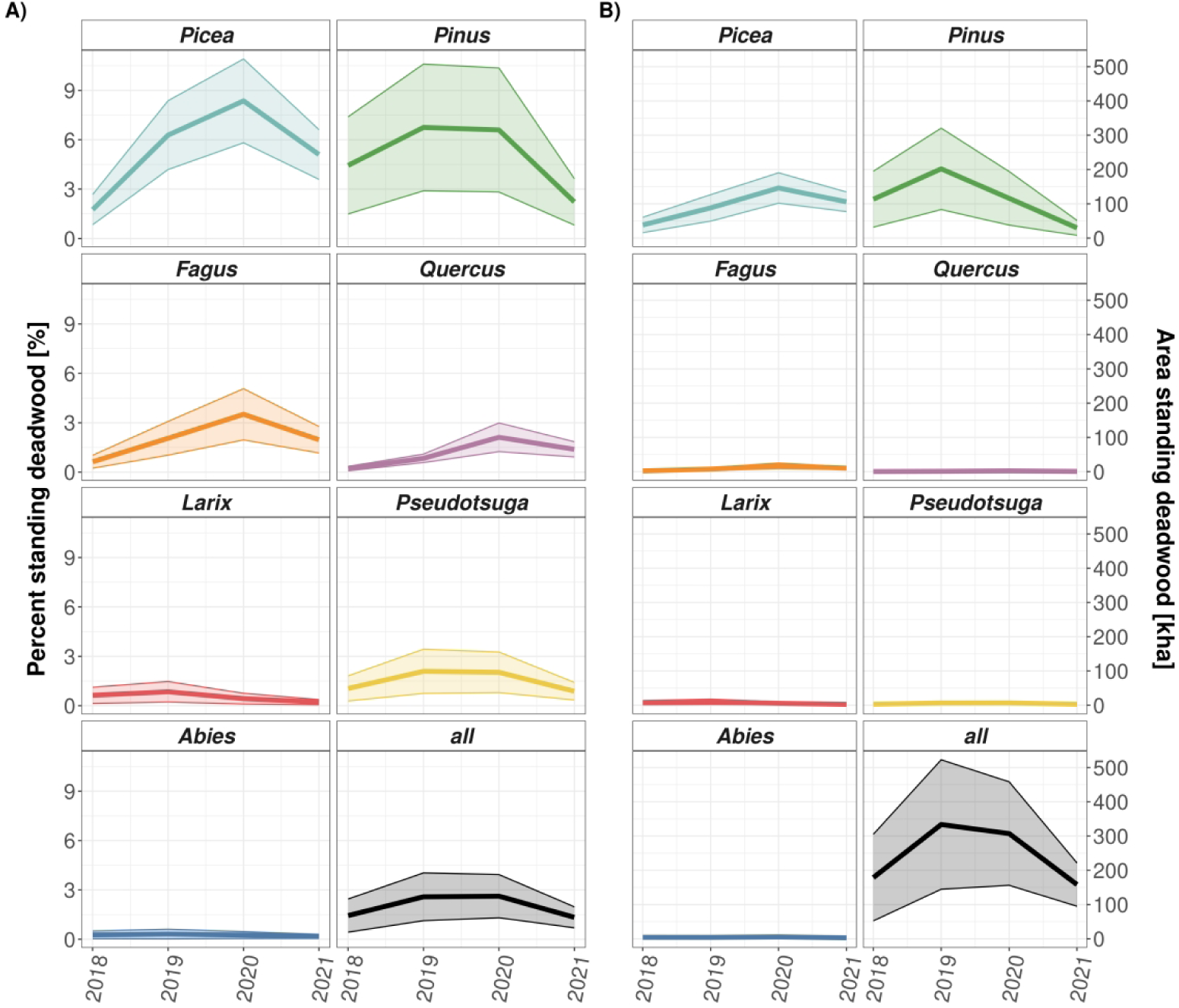
Temporal development of new standing deadwood from 2018 to 2021 for main tree species (*Abies, Fagus, Larix, Picea, Pinus, Pseudotsuga, Quercus*) in Germany. A) The proportion of standing deadwood (% of the respective species), B) total affected area of the species (kha). The thick line represents the mean value of the various threshold values, the correspondingly coloured ribbon the standard deviation.

The mortality patterns between 2018–2021 varied considerably by region (Figure 4a). The most affected regions in terms of area of standing deadwood (Figure 4b) were Süder Uplands (region ID D38, 74.2 kha) and Harz (D37, 47.2 kha), followed by Fläming Heath (D11, 26.5 kha), Elbe-Mulde-Plain (D10, 26.1 kha), Brandenburg Heath and Lake District (D12, 25.1 kha), Lower Saxon Hills (D36, 20 kha), Middle Elbe Plain (D09, 19.3 kha), and Wendland and Altmark (D29, 19 kha). The most affected regions in terms of percentage of standing deadwood (Figure 4c) were Harz (D37, 30.2%), Elbe-Mulde-Plain (D10, 21.4%), Middle Elbe Plain (D09, 21.1%), Saxon-Bohemian Chalk Sandstone Region (D15, 18.4%), Süder Uplands (D38, 17.1%), Wendland and Altmark (D29, 16.8%), Fläming Heath (D11,16.4%), and Lusatian Basin and Spreewald (D08, 12.6%). A detailed overview of the accumulated standing deadwood during 2018–2021 for the major natural regions of Germany can be found in the supplementary materials (Table S3).

**Figure 4.**
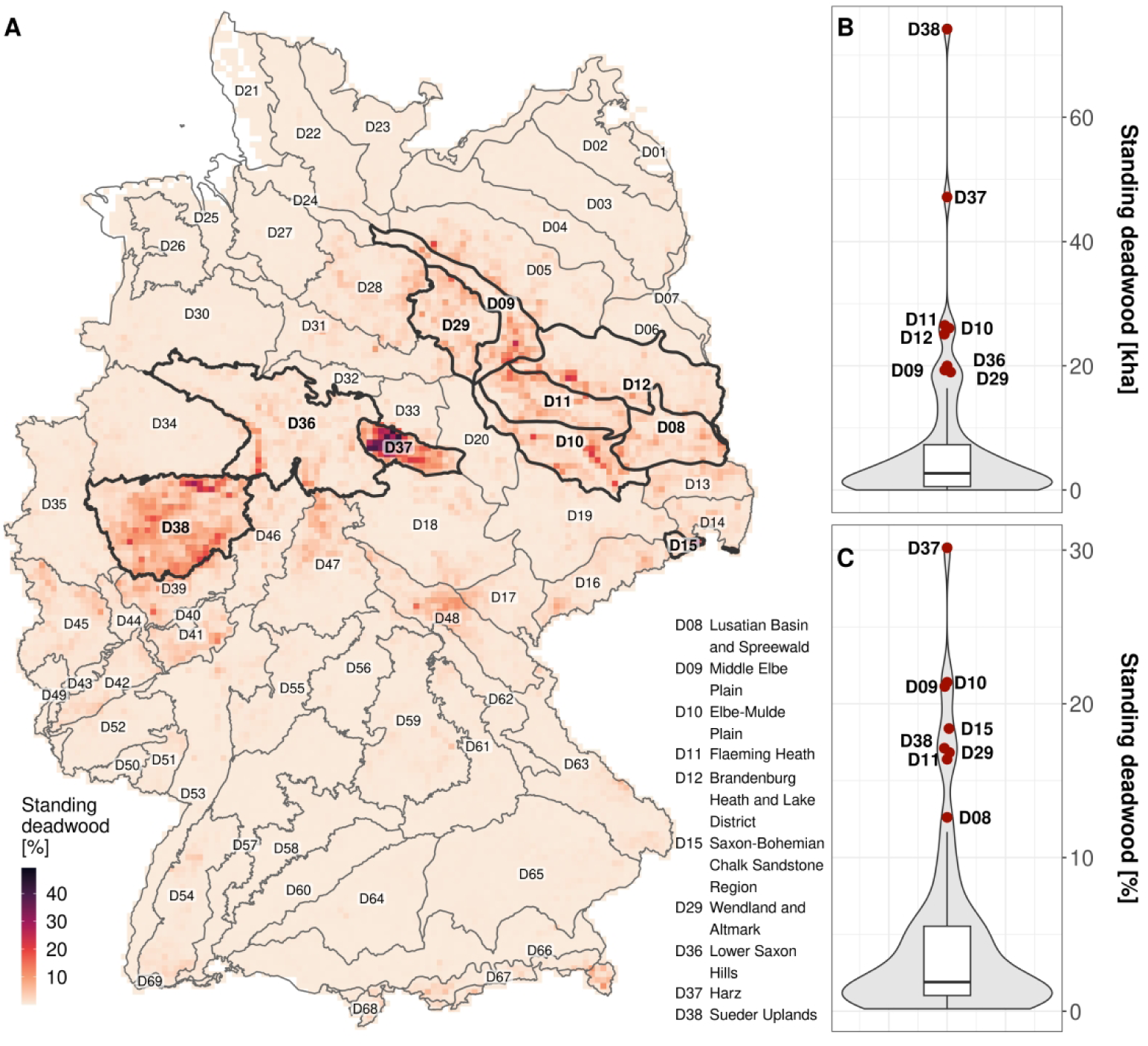
Accumulated standing deadwood from 2018–2021 for the major natural regions in Germany. (A) Map of accumulated standing deadwood in percent of the forested area. For visualisation purposes the maps from (Schiefer et al., 2023) were aggregated to 5 km spatial resolution. Most affected regions are highlighted with a bold outline, and region names along with the IDs (according to the official naming convention) are given in the figure. For the names of all other regions see supplementary materials. (B) Total area [kha] and (C) percent [%] of affected forest areas for the major natural regions in Germany.

The mean explained variance (coefficient of determination R²) of the species-specific random forest models was 0.68 for *Pinus*, 0.64 for *Abies*, 0.6 for *Fagus*, 0.46 for *Quercus*, 0.43 for *Picea*, 0.43 for *Pseudotsuga*, and 0.36 for *Larix*. The random forest permutation importance (Figure 5) reveals that climate and weather were the most important environmental factors explaining standing deadwood dynamics. Late frost was the most important variable (median permutation importance RMSE across species = 15.3%) followed by canopy height (15.2%) as the only non-climate/weather variable in the top group. Variable importances ranked three to ten were climatic water balance (14.3%), hot days (13.7%), climatic water balance of the prior year (13.2%), vapor pressure deficit (12.1%), late frosts of the prior year (11.7%), hot days of the prior year (11.6%), precipitation seasonality (10.6%), and precipitation of the warmest quarter (10.5%). Least important climatic variable was temperature annual range (7.4%, rank 18). The edaphic variables soil drought intensity (9.8%), sand content (9.4%), and soil drought intensity of the prior year (9%) were less important for the models than the climatic variables. Net primary productivity (9.2%) and biomass (9%) ranked 13 and 15 in variable importance. The forest stand-related variables stand age (8.3%) and tree species richness (7.3%) ranked 17 and 19. Slope (9.1%) was the most important variable with the topographic site conditions. The aspect variables northness (4.7%) and eastness (4.1%) were the overall least important variables. *Abies* showed the highest importances for most of the variables, except for canopy height, biomass, stand age, and tree species richness. Lowest importances were observed for *Pinus* for most of the variables. Variables from the year of first of occurrence of standing deadwood were more important than the respective variables from the prior year.

**Figure 5.**
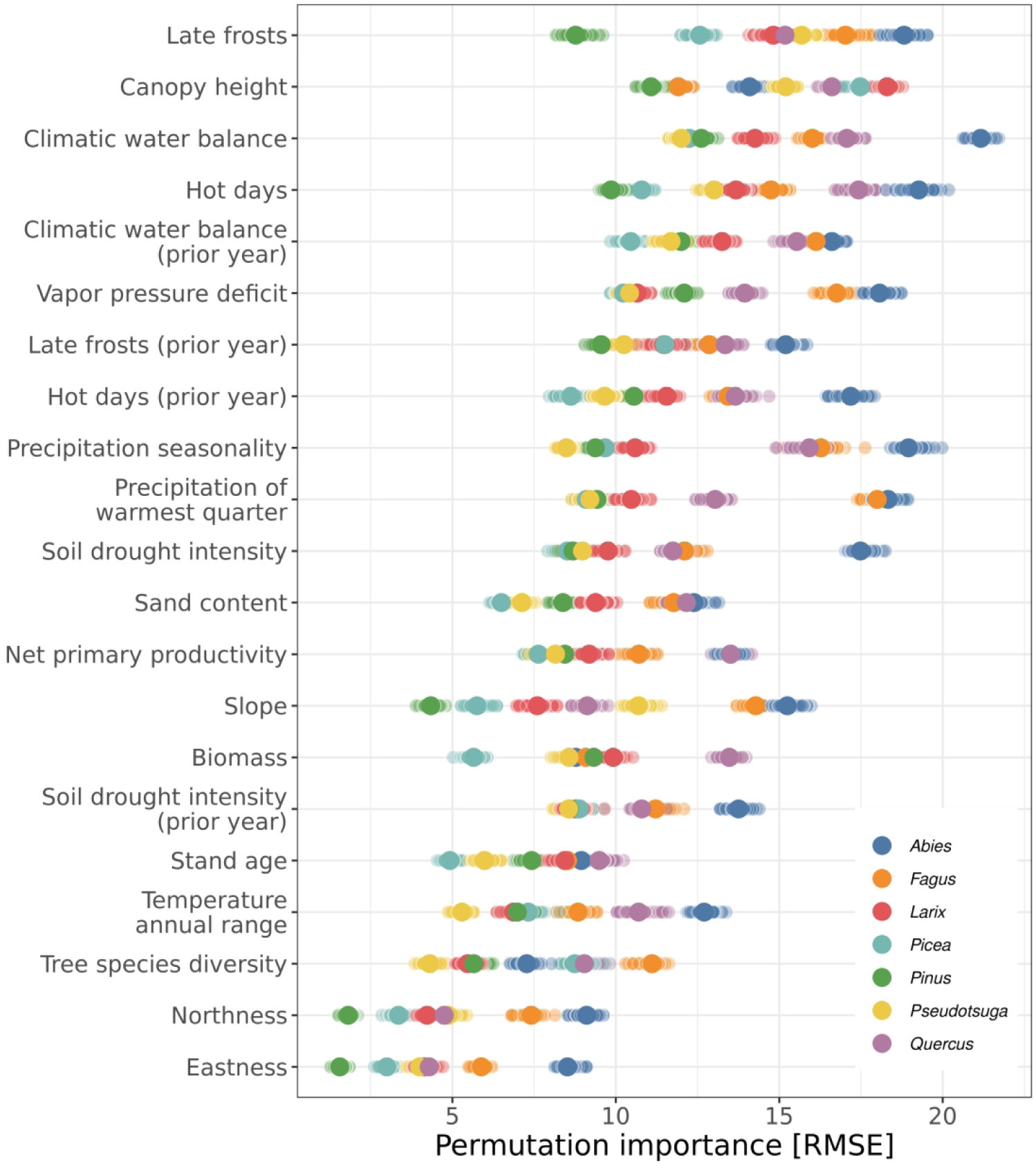
Variable permutation importance of the random forest models. Variables are in descending order according to their median variable importance over all species.

Detailed relationships were obtained using the accumulated local effect (ALE) plots created for each of the 21 predictor variables (Figure 6). Late frosts showed no consistent effect across species despite its high variable importance. *Pseudotsuga*, *Picea*, and *Abies*, however, showed a decreased mortality with more days of late frost. For *Fagus* the effect was reverse with an increased mortality with more days of late frost. Late frosts of the prior year showed a clearer picture with an increased mortality with more days with late frosts for *Quercus*, *Pinus*, *Fagus*, and *Larix*. *Abies*, *Pseudotsuga*, and *Picea* were again more tolerant to late frosts. The effect of canopy height was similar across all species with smaller canopy heights linked to more standing deadwood and taller trees linked to less standing deadwood. This effect was largest for *Larix* and smallest for *Abies*. The effect of the climatic water balance was the same for all species and a more negative climatic water balance was linked to more standing deadwood, except for *Abies* that was inverted. For *Picea*, a smaller deficit was sufficient for increased mortality. The climatic water balance of the prior year showed the same general trends but with a smaller effect. The effect of hot days on the fraction of standing deadwood was similar for all species with higher mortality with more hot days. When exceeding 11 to 17 hot days depending on the species, more standing deadwood was observed. The effect of more hot days on standing deadwood was stronger for the current than the previous year. The largest effect of hot days for both years was observed for *Abies* and *Fagus*.

**Figure 6.**
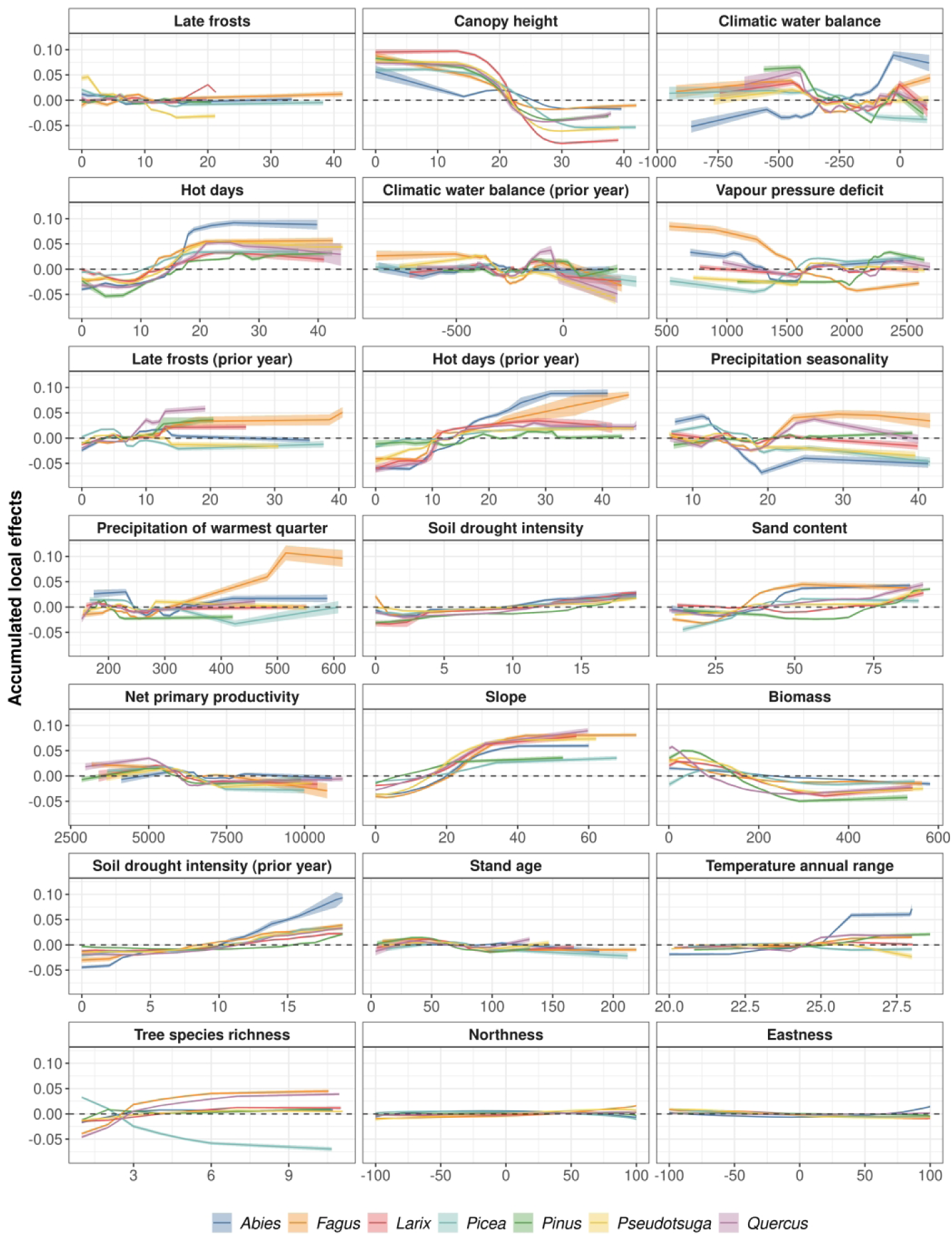
Accumulated local effect (ALE) plots for the 21 environmental predictor variables in descending order based on variable permutation importance. A positive value on the y-axis means contributing to more standing deadwood, negative values to less. ALEs are depicted separately for the seven main tree species (*Abies, Fagus, Larix, Picea, Pinus, Pseudotsuga, Quercus*) in Germany. The solid lines show the mean effect across all repetitions with the transparent ribbon denoting the standard deviation.

The differences in the effects between the species were greatest for vapor pressure deficit. *Quercus, Larix, Pinus, Pseudotsuga,* and *Picea* behaved similarly with increasing fraction of standing deadwood at higher vapor pressure deficits. However, the deficit at which the effect changed to more standing deadwood was considerably higher for *Pinus* at 2.1 kPa compared to 1.5-1.7 kPa for the other four species. For *Abies* and *Fagus* a lower vapor pressure deficit was linked to more standing deadwood and for *Fagus* a higher deficit even was linked to less standing deadwood. The effect of precipitation varied among the species with higher seasonality. *Picea*, *Pseudotsuga*, and *Abies* benefitted from higher precipitation seasonality and standing deadwood decreased, whereas standing deadwood for *Fagus* and *Quercus* increased. We observed increased mortality of *Fagus* in regions with a higher total precipitation during the warmest quarter. Higher soil drought intensities (both years) were linked to increased mortality, but the accumulated local effect was small. Only *Abies* experienced higher mortality rates than the other species when soil drought intensity of the year prior dieback was greater than 10. The species showed different sensitivities towards sand content and its associated low water retention capacity. Whereas standing deadwood only increased at sand contents larger than 75% for *Picea, Quercus, Larix, Pseudotsuga*, and *Pinus*, higher amounts of standing deadwood were already observed at 30% sand content for *Fagus* and *Abies*. Lower net primary productivity and lower biomass were both linked to increased amounts of standing deadwood. Stand age showed only a small effect with slightly increased mortality for younger stands. For the topographical conditions, slopes above 15° featured to more standing deadwood, but the aspect components northness and eastness had no clear effect. For *Larix, Pseudotsuga, Pinus*, and *Abies*, tree species richness had no clear effect. A higher tree species richness was linked to more standing deadwood for *Quercus* and *Fagus*. *Picea* showed a contrasting effect, with less standing deadwood in more diverse forests.

## Discussion

### Pattern of tree mortality in Germany

The 2018–2021 summer droughts resulted in excess tree mortality rates in large parts of Central and northeastern Germany. Compared to the nationwide forest condition survey (BMEL, 2023), both assessments reveal a legacy effect with standing deadwood only increasing sharply in the second consecutive drought year in 2019. However, our results indicate overall higher amounts of standing deadwood and an earlier increase in dieback after the 2018–2021 summer droughts. Whereas in 2018 the statistics from the forest condition survey report 0.51% of standing deadwood in Germany (BMEL, 2023), our analysis resulted in 1.4 ± 1.0%. In 2019 this percentage already increased to 2.5 ± 1.4% (for the single year) in our analysis, but only 0.9% (accumulated) in the survey. The steep increase in standing deadwood in 2020 was also mirrored in the forest condition survey with 2.25%, while our analysis revealed 2.6 ± 1.3% standing deadwood. The highest percentage of standing deadwood in the forest condition survey is reported in 2021 with 2.35% in contrast to the 1.3 ± 0.6% found in our study. Species-specific statistics of standing deadwood from the forest condition monitor accumulated over the same years were considerably lower than found in our study with *Fagus*: 2.4% (this study 8.2 ± 3.8%), *Picea*: 10.4% (21.5 ± 7.1 %), *Pinus*: 3.8% (20 ± 12%), and *Quercus*: 2.8% (4.5 ± 1.7%). We suspect several reasons for the differences between our analysis and the forest condition survey: firstly, the forest condition survey is based on observations of individual dead trees (n = 9688 from 402 sample plots according to BMEL, 2023) and is reported on the total number of trees in the survey. Whereas a large number of small individuals that were not affected by mortality might result in lower estimates of the forest condition survey, larger tree individuals might have an overly huge impact on the standing deadwood cover obtained from the remote sensing perspective. Therefore, a direct comparison of this individual tree-based survey with the area-based remote sensing analysis is hampered. Secondly, the differences may be explained by the different perspectives, as remote sensing assesses tree mortality patterns from the bird’s eye perspective, while the forest condition survey data is recorded from the ground. The upper parts of the canopy are subject to higher levels of solar radiation, increased atmospheric coupling, and long water conduits to the canopy and are, hence, particularly susceptible to dieback. The upward looking perspective of the forest condition survey can impede the view of this upper canopy (e.g., due to crown overlap and dense undergrowth) and result in an underestimation of standing deadwood. Lastly, our estimates of standing deadwood from Schiefer et al. (2023) do not necessarily include only tree individuals but also partially dead tree crown or branches, which may result in higher estimates of standing deadwood.

The temporal patterns of tree mortality obtained in this study mirror those of Thonfeld et al. (2022) and Global Forest Watch (GFW, 2023; Hansen et al., 2013). However, with 978 ± 529 kha from 2018– 2021 our results suggest almost twice the amount of standing deadwood compared with 501 kha mapped by Thonfeld et al. (2022) and 543 kha mapped by the Global Forest Watch (GFW, 2023; Hansen et al., 2013). Although both products do not explicitly map tree mortality but also include other sources of forest loss, for example logging and windthrow, they detect considerably less forest loss. Thonfeld et al. (2022) used Sentinel-2 and Landsat 8 time series and calculated anomalies in the disturbance index. Index based approaches circumvent the need for training data for supervised mapping approaches but cannot explicitly map mortality and therefore become hard to validate and interpret. The coarser spatial resolution of Landsat 8 (30 m) together with a binary classification (dead, not dead) is not an ideal basis to represent the scattered nature of canopy dieback and might, hence, lead to a general underestimation of tree mortality (Cheng et al., 2024). Likewise, the forest loss maps from the Globel Forest Watch (GFW, 2023; Hansen et al., 2013) also rely on the 30 m Landsat data and a binary classification. Accordingly, they cannot robustly represent small-scale tree mortality patterns (see Figure 7) (e.g., single dead tree crowns or patches thereof) and, hence, might underestimate tree mortality (Galiatsatos et al., 2020; Hartmann, Schuldt, et al., 2018). Only 26.6 ± 14.5% of the pixel classified as standing deadwood in Schiefer et al. (2023) were also detected as forest loss by GlobalForestWatch (Hansen et al., 2013). This indicates that a high spatial resolution is required to also capture scattered processes of tree mortality, and that tree mortality in Germany appears to be to be greater than previously expected.

**Figure 7.**
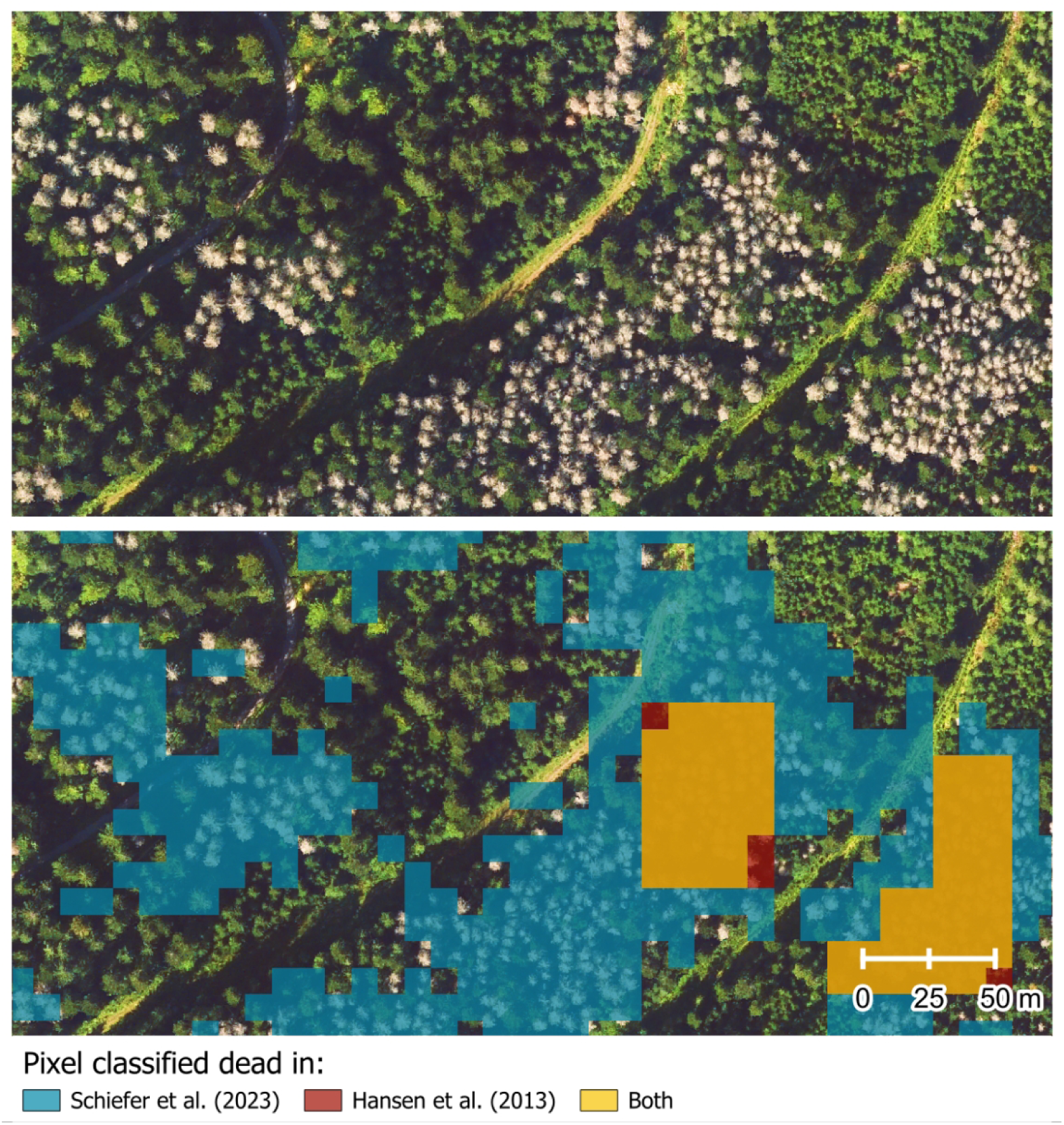
Comparison of the two tree mortality products from Schiefer et al. (2023) and GlobalForestWatch (Hansen et al., 2013). With the spatial resolution of 10 m, Schiefer et al. (2023) also detects scattered occurrences of standing deadwood. For GlobalForestWatch the coarser spatial resolution of 30 m only detects larger patches of deadwood.

Deviations from the standing deadwood cover fractions with other remote sensing products or field-based surveys may also stem from biases of the standing deadwood map used in this study. For example, we observed that sometimes the product of Schiefer et al. (2023) overpredicted tree mortality near forest edges, roads, and paths. Another potential source of overestimation in our study is the sparse canopy of some species, such as *Pinus.* Especially under dry conditions, defoliation and mortality of stands are often observed for *Pinus* (Bose et al., 2024) and the forest floor subsequently showing through the sparse canopy might be interpreted as a standing deadwood signal. Insect infestation can also lead to defoliation (Haynes et al., 2014; Skrzecz et al., 2020), which further thins out the crown, but the trees subsequently recover. This may be particularly true for the *Pinus-* dominated northeastern parts of Germany and is reflected in the large standard deviation of standing deadwood from the sensitivity analysis of the different thresholds (see Figure 3). The respective region was not recorded as being so severely affected by Thonfeld et al. (2022). However, for coniferous species the authors excluded pixels with NDVI values smaller 0.5 from the analysis, which might underestimate standing deadwood in the case of *Pinus*, as the NDVI of *Pinus* is generally not very pronounced. Moreover, the proportion of deadwood reported in the national forest condition survey also increased considerably for *Pinus* (from 1.4% to 2.2% deadwood in 2023) and *Picea* (from 4.5% to 7.4% in 2022). This might indicate a lag effect introduced in the consecutive drought years 2018–2019 and that the proportion of deadwood is in fact higher than previously assumed.

Overall, we found coniferous species to show much higher rates of tree mortality, especially *Pinus* and *Picea* but also *Pseudotsuga*, compared to deciduous species. In the underlying tree species map (Blickensdörfer et al., 2024) *Pseudotsuga* and *Abies* showed the lowest accuracies (User’s accuracy 37.07 ± 1.33 for *Pseudotsuga* and 24.65 ± 1.95 for *Abies*) and were often confused with *Picea.* Therefore, standing deadwood of *Pseudotsuga* and *Abies* is likely to be overestimated in our analysis. In our assessment, conifers often died back over larger areas in a single event, while for deciduous species smaller patches of standing deadwood accumulated over the years. This observation was expected given that the deciduous species usually show gradual die-back of the canopy rather than sudden tree death, by contrast to coniferous species.

### Atmospheric conditions as main predictors of tree mortality

We identified atmospheric conditions (i.e., late frosts, climatic water balance, hot days, and vapor pressure deficit) to be the most important predictors of tree mortality. The most important variable was late frost, although the accumulated local effect was small. One reason for this may be that late frosts tend to have a large effect at a local level and are hence very important, but do not constitute a universal effect across the whole of Germany. Late frost occurs when tree buds have burst and leaves sprouted after spring warming, followed by temperatures well below zero. Late frosts often occur locally which might explain the small accumulated local effects in our large-scale analysis. Whereas a decreasing mortality with more days of late frosts indicate a tolerance to frost for *Pseudotsuga*, *Picea*, and *Abies*, especially for *Quercus* and *Fagus* (but also *Pinus*), more late frosts were associated with more standing deadwood. In the future, it is expected that late frosts will increase as a result of climate change (Vautard et al., 2023) putting vast areas of broadleaved trees at particular risk, given they are more susceptible to late frost events then conifers (Fisichelli et al., 2014). Hence, air temperatures are an important environmental variable, not only because of temperature extremes (i.e., late frost and hot days) but also because of the relatively important vapor pressure deficit which is largely driven by temperature. The only temperature related variable that was less important was temperature annual range, which in line with previous studies indicates that differences in the temperature range are less decisive than temperature extremes (Adams et al., 2009; Anderegg et al., 2013).

As generally expected, we found a more negative climatic water balance to increase tree mortality for all species, except for *Abies,* which showed an inverted pattern. Since *Abies* largely occurs at higher altitudes with higher precipitation, the precipitation amounts might be still sufficient to compensate for the observed effect of the climatic water balance.

Our results on the importance of the respective environmental variables (see Figure 5) confirm the importance of hotter droughts, in which atmospheric drought is accompanied by heat (Allen et al., 2015; Hartmann et al., 2022), for tree mortality during 2018–2020. Soil conditions were not as important as atmospheric drought and high temperatures. During drought, some tree species are able to compensate for water loss and prevent the death of plant material through various mechanisms (McDowell et al., 2022). Under simultaneous heat and drought, i.e., hotter droughts, negative impacts are aggravated because of greater atmospheric demands (higher vapor pressure deficit) and greater soil moisture limitations (Grossiord et al., 2020; Hartmann et al., 2022). In the first drought year in 2018, the trees already suffered damages (Schuldt et al., 2020) and thus started with significant abiotic and biotic legacies, such as depleted soil water reservoirs and damages to the tree’s water transport system (McDowell et al., 2022), into the consecutive drought years 2019 and 2020. These legacies likely induced the pronounced increases in tree mortality we observed during the subsequent drought years. Similarly, tree growth and physiological stress responses were found to be even more pronounced in 2019 than in 2018 as result of such abiotic and biotic legacies (Schnabel et al., 2022). The predicted increase in such consecutive and hotter droughts (Hari et al., 2020) will likely further increase the risk for forest dieback events in the future.

### Mixed effect of forest composition and structure

#### Canopy height

Canopy height was the second most important variable based on random forest permutation importances, with a consistent effect across all species that smaller canopy heights showed greater mortality and were more affected than tall canopies. At first glance, these findings my contradict results from studies obtained on a local level, where tree height largely explained mortality risks during droughts leaving especially larger and older trees at risk due to hydraulic vulnerability and ensuing pest infestations (Bennett et al., 2015; Gora & Esquivel-Muelbert, 2021; Stovall et al., 2019, 2020). However, canopy height and tree height are not directly comparable. Canopy height represents the average height of an area (here a 10 m pixel) and thereby also aggregates the overall canopy cover. Accordingly, a low canopy height can result from small trees or gaps in a canopy. Thus, our results indicate that open forest stands as well as forest edges may be particularly prone to tree mortality. This importance of the canopy structure and edge effects for drought-induced tree mortality were highlighted by previous studies and explained by reduced climatic buffering capacities of open canopies stands or forest edges (Anderegg et al., 2019; Buras et al., 2018; Jönsson et al., 2007). Thus, our findings at the landscape-scale support the findings of Stephenson & Das (2020), who found mortality to be very dependent on height-related changes in forest composition and not only governed by an intrinsically higher vulnerability of higher trees.

#### Site productivity and biomass

With increasing atmospheric CO_2_ concentrations, increasing average temperatures and a longer growing season in the future an increase in forest productivity is expected (Lindner et al., 2014; N. G. McDowell et al., 2020). Higher site productivity increases growth rates of trees and may induce a higher susceptibility to embolisms and hydraulic failure during drought conditions (Pretzsch et al., 2018). At the same time, older trees usually have a reduced plasticity to changing environmental conditions and higher biomass that comes with higher maintenance costs reducing soil water availability faster and therefore increase drought stress and mortality risk (Jump et al., 2017). In contrast, our results showed lower mortality rates in stands with a higher net primary productivity and a larger biomass. This is in line with studies at the stand level, that found older forests to be more resistant to droughts as these are often more intact forests with closed canopies and enhanced climate buffering capacities (Anderegg et al., 2013; N. McDowell et al., 2008; Thom et al., 2019).Thus, it appears that relationships to stand age and productivity are complex and hard to generalize. There might be several confounding factors such as the soil substrate or the rooting depths of species, which determine the plant available water during drought and their strong control on drought resilience (Brunner et al., 2015; Trugman et al., 2021).

#### Tree species and species richness

As expected, we observed strong differences in tree mortality patterns across species and their relationships with environmental variables (Figures 5 & 6). These variations can largely be attributed to the differing drought resistance of tree species, with some being more resilient to climate extremes or climatic changes than others (Schuldt et al., 2020). While the effect of species was very pronounced, we did not find a clear relationship of forest mortality with tree species richness. Overall, the effect of tree species richness on tree mortality remains debated. Former studies on tree species richness effects on mortality yielded mixed results, with some species profiting from diversity and others not depending on their functional traits (Blondeel et al., 2024; Depauw et al., 2024; Liu et al., 2022; Searle et al., 2022). For instance, Hajek et al. (2023) reported both synergistic tree species interactions in mixtures depending on the examined species during the 2018 hotter drought in an experimental setup. Across Germany, a higher degree of conspecific neighbours was recently shown to increase mortality risks relative to heterospecific neighbours using national forest inventory data in a study by Kulha et al. (2023). Here, we found that a higher tree species richness was not generally linked to lower tree mortality for most species and that tree species richness was generally less important than abiotic predictors and canopy height for explaining mortality patterns. When interpreting the results, it should be considered that the tree species map used here shows the main tree species per pixel and that a mixture with other tree species cannot be ruled out. The two broadleaved species *Fagus* and *Quercus* experienced higher mortality with increased tree species richness. One may speculate that the increased tree mortality for these species in mixed forests is caused by a higher competition for water in more productive mixed stands (Bauhus et al., 2017). However, our purely observational approach does not allow testing this assumption or rule out a potential confounding of this diversity effect with other abiotic variables, which would require explicit comparisons of mixtures with all of their constituent monocultures under comparable site conditions (see Baeten et al., 2013; Depauw et al., 2024). Therefore, future studies should aim to decipher the mechanisms that caused the increased mortality rates of *Fagus* and *Quercus* we observed. In contrast, *Picea* benefited from more diverse forest stands and we observed larger amounts of standing deadwood in monocultures. There is ample evidence that species mixing reduces the effect of specialist pests and pathogens (Jactel et al., 2021; Messier et al., 2022). Especially for *Picea* the reason for large-scale die-off is therefore the interaction of drought stress and bark beetle outbreaks, which can only propagate at this extent in vast monocultural stands. *Pinus* is also often planted in monocultures in Germany and died over extensive areas during 2018–2021. In contrast to *Picea*, however, the diversity of tree species was not related to mortality, presumably due to the regional lack of specialized pests. Stephenson et al. (2019) also highlight this effect and found the combined effects of the presence of specialized bark beetles and drought stress to greatly increase tree mortality. Despite the current absence of specialized pathogens and pests for some tree species, under changing climatic conditions new specialized diseases are likely to spread and to put currently less affected tree species at risk.

### Conclusion

Using a wall-to-wall remote sensing product with high spatial detail, we revealed almost twice as much standing deadwood in Germany during 2018–2021 than other ground-based inventories or previous remote sensing assessments. We attribute these large differences mainly to i) the increased coverage and spatial detail of our approach, ii) the fundamental differences of ground-based sampling protocols and the bird perspective from Earth observation satellites. Overall, remote sensing provides a valuable landscape-level view on tree mortality, complementing the insights on tree mortality processes we have from ground-based forest assessments. We found a complex interplay of environmental predictors, with extreme atmospheric conditions, i.e. hotter droughts but also late frosts, being the most important predictors for the observed tree mortality. The species’ response varied greatly, and the revealed patterns provide important information for climate change adaptation. Mainly the coniferous species *Picea* and *Pinus* died as they appear to be not well adapted to hotter droughts. However, the extreme atmospheric conditions also put broadleaved species at risk, and particularly late frost events after vegetation onset played an important role in this respect. Our results showed that stand structure and composition influences tree mortality and, hence, that management of forest structure and composition can help mitigate the risks of global warming. Our results at the landscape level suggest that small, young stands contributed to more overall tree mortality in Germany. Species information was crucial for interpreting patterns and predictors of tree mortality underlying the need for species-specific databases for upscaling local observations of standing deadwood to the landscape level and elucidating underlying drivers. To gain insights on tree mortality processes at continental or global scales adequate maps of standing deadwood are required. This can only be accomplished if the scientific community contributes to the collective gathering of reference data from multiple regions and biomes to improve the upscaling of standing deadwood observations from local-level to landscape-level, e.g. in initiatives such as www.deadtreath.earth. Due to limitations in computing power and models, environmental variables are often only considered as observations of single points in time. The provision of corresponding time series data and the development of suitable models for investigating the complex spatiotemporal patterns of tree mortality should be advanced in view of the global increase in tree mortality.

## CRediT author contributions

**Felix Schiefer**: Conceptualization, Methodology, Software, Validation, Formal analysis, Data curation, Writing—Original Draft, Visualization. **Sebastian Schmidtlein**: Supervision, Resources, Writing— Review & Editing. **Henrik Hartmann**: Writing—Original Draft, Writing—Review & Editing. **Florian Schnabel**: Writing—Original Draft, Writing—Review & Editing. **Teja Kattenborn**: Conceptualization, Methodology, Supervision, Writing—Original Draft, Writing—Review & Editing, Funding acquisition.

## Data availability statement

The data underlying this article are available in the KITopen repository, at https://dx.doi.org/10.5445/IR/1000158765. Code for the conducted analyses is available on GitHub, at https://github.com/FelixSchiefer/TreeMortalityPattern.

## Supplementary material

Supplementary data is available at *Forestry* online.

