## Supplementary Material for "Large-scale remote sensing reveals that tree mortality in Germany appears to be greater than previously expected"

**Table S1:** Originally selected environmental predictor variables

| Variable | Spatial resolution | Unit | Rationale (include/*exclude*) | Data source |
|---|---|---|---|---|
| Slope | 10 m | ° | Topographic site conditions | Slope and eastness (sinus) and northness (cosine) of terrain aspect derived from Copernicus DEM EEA-10 (2021) |
| Eastness | | *rad* | | |
| Northness | | *rad* | | |
| Sand content | 30 m | *%* | Soil type | Hengl, T. and Parente, L. (2022) |
| Stand age | 100 m | *years* | Hypothesis that older trees are more susceptible to mortality (cf. Socha et al., 2023) | Besnard et al. (2021) |
| Canopy height | 10 m | *m* | Hypothesis that taller trees are more susceptible to mortality (cf. Stovall et al., 2019, 2020; Bennett et al., 2015) | Lang et al. (2023) |
| Tree species richness | 10 m | *no.* | Hypothesis that tree species richness mitigates mortality (cf. Anderegg et al., 2018; Schnabel et al., 2021; Depauw et al., 2024; Grossiord et al., 2020; Searle et al., 2022) | Calculated from Blickensdörfer et al. (2024) tree species map as number of tree species in a 100 m buffer |
| Mean diurnal temperature range | 30 sec | *°C* | Climatic site conditions; *Correlated with temperature annual range that better reflects the average site conditions* | Derived from Fick & Hijmans (2017) |
| Temperature annual range | | *°C* | Climatic site conditions | |
| Mean temperature of warmest quarter | | *°C* | Climatic site conditions; *Correlated with vapor pressure deficit* | |
| Annual precipitation | | *mm* | Climatic site conditions; *Correlated with precipitation of warmest quarter which is more meaningful for vegetation growth* | |
| Precipitation seasonality | | *mm* | Climatic site conditions | |
| Precipitation of warmest quarter | | *mm* | Climatic site conditions | |
| Late frosts* | 1 km | *days* | Frost damage after bud burst (cf. Vanoni et al., 2016) | Calculated from DWD Climate Data Center (CDC) (2023b) and (Schimanke et al., 2021) |
| Soil drought intensity* | 4 km | - | Edaphic site conditions | Zink et al. (2016); during vegetation period |
| Total precipitation* | 1 km | *mm* | Climatic site conditions; *Correlated with vapor pressure deficit that incorporates both* | Sum over growing period (March-October); DWD Climate Data Center (CDC) (2023d) |
| Mean Temperature* | 1 km | *°C* | Climatic site conditions; *Correlated with vapor pressure deficit that incorporates both* | DWD Climate Data Center (CDC) (2023c) |
| Vapor pressure deficit | 5.5 km | *Pa* | Driving hydraulic failure | Calculated as 75%-quartile of daily maximum during growing period (March-October) from reanalysis data (Schimanke et al., 2021) |
| Climatic water balance* | 1 km | *mm* | Decreasing water availability increases mortality risk | Calculated from DWD Climate Data Center (CDC) (2023d, 2023e) as anomaly from DWD Climate Data Center (CDC) (2023f) |
| Hot days* | 1 km | *days* | Heat-induced tree mortality through cavitation of water columns within the xylem (cf. Allen et al., 2010) | DWD Climate Data Center (CDC) (2023a) |
| Biomass | 100 m | *Mg/ha* | Proxy for site productivity. Hypothesis that higher productivity enhances susceptibility to drought-induced mortality (cf. Socha et al., 2023) | ESA Biomass CCI (Santoro & Cartus, 2023) |
| Net Primary Productivity | 500 m | *kgC/m²/ year* | Proxy for site productivity. Hypothesis that higher productivity enhances susceptibility to drought-induced mortality (cf. Socha et al., 2023) | Running, S. and Zhao, M. (2021) |

*Variable also for the year prior to first occurrence of standing deadwood

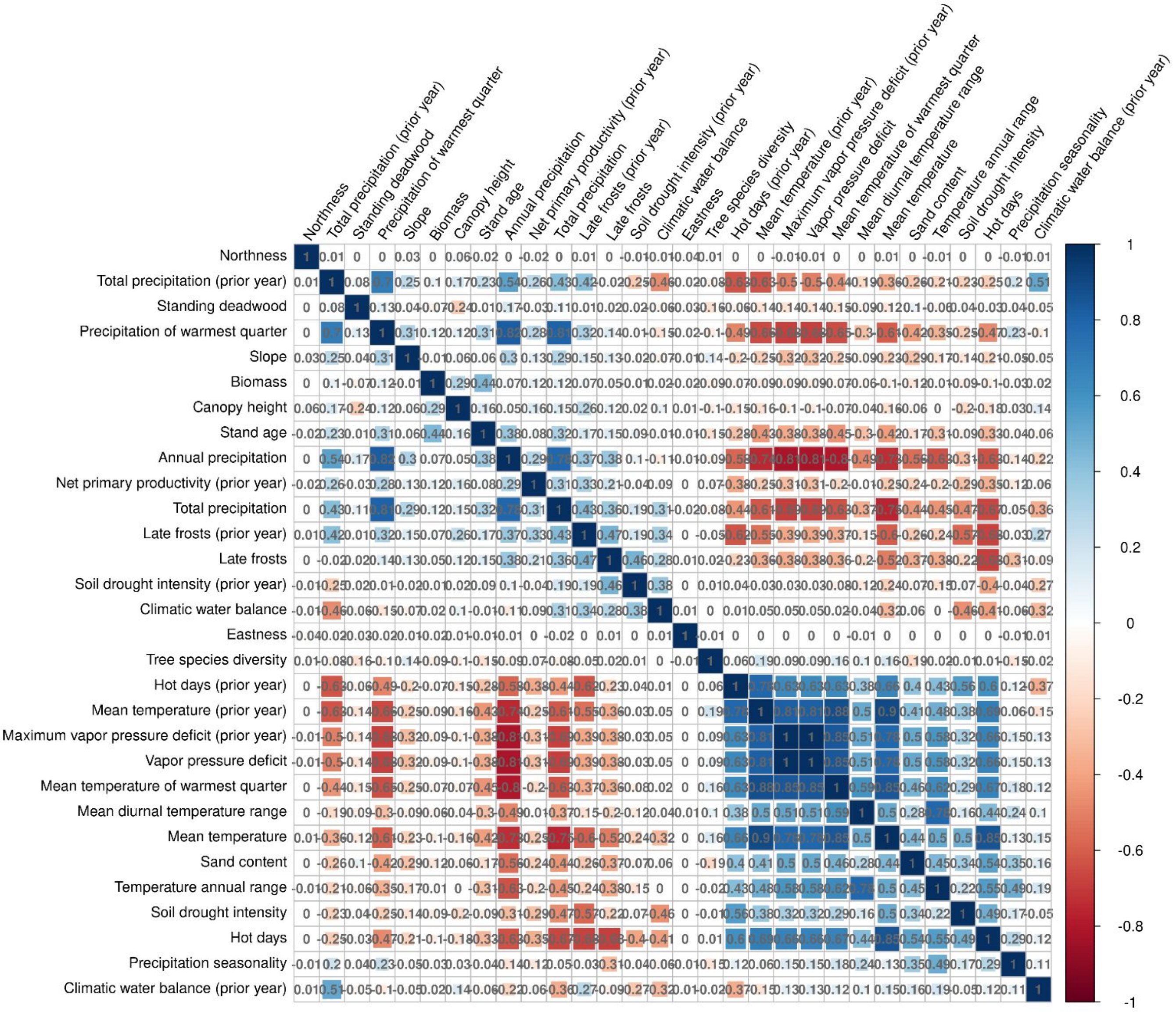


**Figure S1:** Correlation plot for all environmental predictor variables and standing deadwood. Variables with correlations higher than 0.7 were dropped for the analysis.

**Table S2**: Standing deadwood for the seven dominant tree species in Germany over the years 2018–2021

| Species | Year | Area [kha] (mean) | Area [kha] (sd) | Percent [%] (mean) | Percent [%] (sd) |
|---|---|---|---|---|---|
| all | 2018 | 178.67 | 126.26 | 1.44 | 1.01 |
| | 2019 | 333.68 | 188.89 | 2.58 | 1.45 |
| | 2020 | 307.24 | 151.05 | 2.62 | 1.31 |
| | 2021 | 158.27 | 63.14 | 1.33 | 0.64 |
| | acc. | 977.87 | 529.35 | 7.97 | 4.41 |
| *Abies* | 2018 | 4.48 | 4.32 | 0.27 | 0.23 |
| | 2019 | 4.34 | 3.92 | 0.32 | 0.29 |
| | 2020 | 5.71 | 4.81 | 0.26 | 0.21 |
| | 2021 | 3.04 | 2.15 | 0.18 | 0.12 |
| | acc. | 17.57 | 15.20 | 1.03 | 0.86 |
| *Fagus* | 2018 | 2.11 | 1.75 | 0.63 | 0.39 |
| | 2019 | 7.49 | 4.36 | 2.05 | 1.03 |
| | 2020 | 17.09 | 8.27 | 3.51 | 1.56 |
| | 2021 | 10.16 | 4.17 | 1.96 | 0.80 |
| | acc. | 36.85 | 18.55 | 8.15 | 3.78 |
| *Larix* | 2018 | 7.50 | 6.10 | 0.63 | 0.49 |
| | 2019 | 10.10 | 7.81 | 0.84 | 0.62 |
| | 2020 | 5.61 | 4.41 | 0.43 | 0.33 |
| | 2021 | 2.60 | 1.93 | 0.21 | 0.16 |
| | acc. | 25.81 | 20.25 | 2.11 | 1.60 |
| *Picea* | 2018 | 38.16 | 22.38 | 1.75 | 0.92 |
| | 2019 | 88.03 | 38.63 | 6.28 | 2.09 |
| | 2020 | 146.09 | 44.27 | 8.36 | 2.55 |
| | 2021 | 105.79 | 28.65 | 5.10 | 1.51 |
| | acc. | 378.07 | 133.93 | 21.49 | 7.06 |
| *Pinus* | 2018 | 113.30 | 81.60 | 4.44 | 2.95 |
| | 2019 | 201.88 | 118.79 | 6.75 | 3.85 |
| | 2020 | 115.95 | 78.18 | 6.59 | 3.77 |
| | 2021 | 29.58 | 21.77 | 2.22 | 1.41 |
| | acc. | 460.71 | 300.35 | 20.00 | 11.98 |
| *Pseudotsuga* | 2018 | 3.10 | 2.32 | 1.05 | 0.76 |
| | 2019 | 6.54 | 4.43 | 2.09 | 1.34 |
| | 2020 | 6.75 | 4.22 | 2.03 | 1.23 |
| | 2021 | 2.94 | 1.80 | 0.88 | 0.53 |
| | acc. | 19.33 | 12.77 | 6.04 | 3.87 |
| *Quercus* | 2018 | 0.37 | 0.34 | 0.21 | 0.14 |
| | 2019 | 1.00 | 0.61 | 0.84 | 0.26 |
| | 2020 | 2.19 | 1.28 | 2.11 | 0.87 |
| | 2021 | 0.71 | 0.33 | 1.38 | 0.46 |
| | acc. | 4.26 | 2.56 | 4.54 | 1.73 |
| other | 2018 | 9.65 | 7.51 | 1.56 | 1.20 |
| | 2019 | 14.31 | 10.50 | 2.05 | 1.47 |
| | 2020 | 7.86 | 5.78 | 1.31 | 0.95 |
| | 2021 | 3.45 | 2.64 | 0.66 | 0.49 |
| | acc. | 35.27 | 26.43 | 5.57 | 4.12 |

**Table S3:** Accumulated standing deadwood during 2018–2021 for the major natural regions of Germany

| Major natural regions | | | Forest area | Standing deadwood | |
|---|---|---|---|---|---|
| ID | Name (eng.) | Name (ger.) | [kha] | Area [kha] | Percent [%] |
| D01 | Mecklenburg-Western Pomeranian Littoral | Mecklenburgisch-Vorpommersches Küstengebiet | 50.74 | 0.99 | 1.96 |
| D02 | Northeast Mecklenburg Plain and Szczecin Lagoon | Nordostmecklenburgisches Tiefland mit Oderhaffgebiet | 96.15 | 2.98 | 3.10 |
| D03 | Hinterland of the Mecklenburg Lake Plateau | Rückland der Mecklenburg-Brandenburgischen Seenplatte | 155.76 | 5.36 | 3.44 |
| D04 | Mecklenburg Lake Plateau | Mecklenburgische Seenplatte | 265.72 | 9.09 | 3.42 |
| D05 | Mecklenburg-Brandenburg Plateau and Upland | Mecklenburg-Brandenburgisches Platten- und Hügelland | 264.20 | 43.65 | 16.52 |
| D06 | East Brandenburg Plateau | Ostbrandenburgische Platte | 84.71 | 3.52 | 4.16 |
| D07 | Oder Valley | Odertal | 14.16 | 1.34 | 9.43 |
| D08 | Lusatian Basin and Spreewald | Spreewald und Lausitzer Becken- und Heideland | 143.40 | 43.19 | 30.12 |
| D09 | Middle Elbe Plain | Elbtalniederung | 91.51 | 41.32 | 45.15 |
| D10 | Elbe-Mulde Plain | Elbe-Mulde-Tiefland | 121.91 | 54.89 | 45.02 |
| D11 | Fläming Heath | Fläming | 162.09 | 61.35 | 37.85 |
| D12 | Brandenburg Heath and Lake District | Brandenburgisches Heide- und Seengebiet | 322.49 | 61.37 | 19.03 |
| D13 | Upper Lusatian Plateau | Oberlausitzer Heideland | 119.38 | 20.63 | 17.28 |
| D14 | Upper Lusatian | Oberlausitz | 44.92 | 7.38 | 16.44 |
| D15 | Saxon-Bohemian Chalk Sandstone Region | Sächsisch-Böhmisches Kreidesandsteingebiet | 24.69 | 6.96 | 28.20 |
| D16 | Ore Mountains | Erzgebirge | 156.22 | 14.08 | 9.01 |
| D17 | Vogtland | Vogtland | 94.90 | 10.39 | 10.94 |
| D18 | Thuringian Basin and Peripheral Uplands | Thüringer Becken und Randplatten | 220.32 | 21.16 | 9.61 |
| D19 | Saxon Lowland and Saxon Uplands | Erzgebirgsvorland und Sächsisches Hügelland | 96.06 | 11.85 | 12.33 |
| D20 | Eastern Harz Foreland | Mitteldeutsches Schwarzerdegebiet | 11.81 | 3.06 | 25.95 |
| D21 | Schleswig-Holstein Marsh | Schleswig-Holsteinische Marschen und Nordseeinseln | 1.43 | 0.02 | 1.12 |
| D22 | Schleswig-Holstein Geest | Schleswig-Holsteinische Geest | 73.77 | 1.00 | 1.36 |
| D23 | Schleswig-Holstein Uplands | Schleswig-Holsteinisches Hügelland | 68.95 | 0.83 | 1.20 |
| D24 | Lower Elbe Marsh | Unterelbeniederung (Elbmarsch) | 4.10 | 0.19 | 4.69 |
| D25 | Lower Ems and Weser Marshes | Ems-Weser-Marsch | 3.59 | 0.08 | 2.20 |
| D26 | East Frisian Geest | Ostfriesisch-Oldenburgische Geest | 29.19 | 0.45 | 1.54 |
| D27 | Stade Geest | Stader Geest | 84.53 | 1.73 | 2.04 |
| D28 | Lüneburg Heath | Lüneburger Heide | 283.31 | 39.04 | 13.78 |
| D29 | Wendland and Altmark | Wendland und Altmark | 112.86 | 43.94 | 38.93 |
| D30 | Dümmer and Ems-Hunte Geest | Dümmer Geestniederung und Ems-Hunte-Geest | 148.57 | 5.58 | 3.76 |
| D31 | Weser-Aller Plains and Geest | Weser-Aller-Tiefland | 135.62 | 13.90 | 10.25 |
| D32 | Lower Saxony Börde | Niedersächsische Börden | 9.83 | 0.29 | 2.93 |
| D33 | North Harz Foreland | Nördliches Harzvorland | 33.56 | 3.92 | 11.68 |
| D34 | Westphalian Lowland | Westfälische Tieflandsbucht | 122.38 | 6.34 | 5.18 |
| D35 | Lower Rhine Plain and Cologne Lowland | Kölner Bucht und Niederrheinisches Tiefland | 95.55 | 5.65 | 5.92 |
| D36 | Lower Saxon Hills | Niedersächsisches Bergland | 361.52 | 32.05 | 8.86 |
| D37 | Harz | Harz | 156.42 | 65.89 | 42.13 |
| D38 | Süder Uplands | Sauerland | 433.63 | 106.34 | 24.52 |
| D39 | Westerwald | Westerwald | 152.77 | 18.90 | 12.37 |
| D40 | Lahn Valley | Lahntal und Limburger Becken | 9.51 | 0.59 | 6.19 |
| D41 | Taunus | Taunus | 129.17 | 11.42 | 8.84 |
| D42 | Hunsrück | Hunsrück | 153.50 | 8.63 | 5.62 |
| D43 | Moselle Valley | Moseltal | 25.88 | 0.79 | 3.05 |
| D44 | Middle Rhine Valley | Mittelrheingebiet | 35.90 | 2.42 | 6.75 |
| D45 | Eifel and Venn Foreland | Eifel und Vennvorland | 239.97 | 19.88 | 8.28 |
| D46 | West Hesse Uplands | Westhessisches Berg- und Beckenland | 144.00 | 10.41 | 7.23 |
| D47 | East Hesse Upland | Osthessisches Bergland | 326.85 | 21.95 | 6.72 |
| D48 | Thuringian-Franconian Highlands | Thüringisch-Fränkisches Mittelgebirge | 296.61 | 31.41 | 10.59 |
| D49 | Gutland (Bitburg Land) | Gutland (Bitburger Land) | 24.36 | 0.78 | 3.19 |
| D50 | Palatine-Saarland Muschelkalk Region | Pfälzisch-Saarländisches Muschelkalkgebiet | 33.71 | 0.70 | 2.08 |
| D51 | Palatine Forest (the Haardt) | Pfälzer Wald (Haardtgebirge) | 133.20 | 5.79 | 4.35 |
| D52 | Saar-Nahe Hills | Saar-Nahe-Berg- und Hügelland | 158.00 | 3.97 | 2.51 |
| D53 | Upper Rhine Plain | Oberrheinisches Tiefland | 186.21 | 8.12 | 4.36 |
| D54 | Black Forest | Schwarzwald | 397.77 | 10.79 | 2.71 |
| D55 | Odenwald-Spessart-Rhön | Odenwald, Spessart und Südrhön | 339.94 | 6.89 | 2.03 |

| Major natural regions | | | Forest area | Standing deadwood | |
|---|---|---|---|---|---|
| ID | Name (eng.) | Name (ger.) | [kha] | Area [kha] | Percent [%] |
| D56 | Main Franconia Plateau | Mainfränkische Platten | 120.47 | 4.09 | 3.39 |
| D57 | Neckar and Tauber Gäu Plateaus | Neckar- und Tauberland, Gäuplatten | 218.28 | 2.26 | 1.04 |
| D58 | Swabian Keuper-Lias Lands | Schwäbisches Keuper-Liasland | 200.12 | 0.99 | 0.50 |
| D59 | Franconian Keuper-Lias Lands | Fränkisches Keuper-Liasland | 325.97 | 13.29 | 4.08 |
| D60 | Swabian Jura | Schwäbische Alb | 234.05 | 0.84 | 0.36 |
| D61 | Franconian Jura | Fränkische Alb | 281.11 | 4.48 | 1.59 |
| D62 | Upper Palatine-Upper Main Hills | Oberpfälzisch-Obermainisches Hügelland | 117.89 | 3.78 | 3.21 |
| D63 | Upper Palatine-Bavarian Forest | Oberpfälzer und Bayerischer Wald | 347.09 | 10.55 | 3.04 |
| D64 | Iller-Lech Plateau | Donau-Iller-Lech-Platten | 203.23 | 1.41 | 0.69 |
| D65 | Lower Bavarian Upland and Isar-Inn Gravel Plateau | Unterbayerisches Hügelland und Isar-Inn-Schotterplatten | 288.57 | 4.38 | 1.52 |
| D66 | Pre-Alpine Hills and Moorland | Voralpines Hügel- und Moorland | 280.54 | 1.70 | 0.60 |
| D67 | Swabian-Bavarian Pre-alps | Schwäbisch-Oberbayerische Voralpen | 158.87 | 7.37 | 4.64 |
| D68 | Northern Limestone Alps | Nördliche Kalkalpen | 50.82 | 7.39 | 14.54 |
| D69 | Dinkelberg and Upper Rhine Valley | Hochrheingebiet und Dinkelberg | 8.79 | 0.10 | 1.19 |
| D70 | German Bight | Deutsche Bucht | - | - | - |
| D71 | Dogger Bank | Doggerbank und angrenzende zentrale Nordsee | - | - | - |
| D72 | Western Baltic | Westliche Ostsee | 0.23 | 0.00 | 1.45 |
| D73 | Eastern Baltic | Östliche Ostsee | 0.76 | 0.03 | 3.90 |
